# Post-transcriptional regulation of human endogenous retroviruses by RNA-Binding Motif Protein 4, RBM4

**DOI:** 10.1101/2020.03.30.017111

**Authors:** Amir K. Foroushani, Bryan Chim, Madeline Wong, Andre Rastegar, Kent Barbian, Craig Martens, Markus Hafner, Stefan A. Muljo

## Abstract

The human genome encodes for over 1,500 RNA-binding proteins (RBPs), which coordinate regulatory events on RNA transcripts (Gerstberger *et al*., 2014). Most studies of RBPs concentrate on their action on mRNAs that encode protein, which constitute a minority of the transcriptome. A widely neglected subset of our transcriptome derives from integrated retroviral elements termed endogenous retroviruses (ERVs) that comprise ~8% of the human genome. Some ERVs have been shown to be transcribed under physiological and pathological conditions suggesting that sophisticated regulatory mechanisms to coordinate and prevent their ectopic expression exist. However, it is unknown whether RBPs and ERV transcripts directly interact to provide a post-transcriptional layer of regulation. Here, we implemented a computational pipeline to determine the correlation of expression between individual RBPs and ERVs from single-cell or bulk RNA sequencing data. One of our top candidates for an RBP negatively regulating ERV expression was RNA-Binding Motif Protein 4 (RBM4). We used PAR-CLIP to demonstrate that RBM4 indeed bound ERV transcripts at CGG consensus elements. Loss of RBM4 resulted in elevated transcript level of bound ERVs of the HERV-K and -H families, as well as increased expression of HERV-K envelope protein. We pinpointed RBM4 regulation of HERV-K to a CGG-containing element that is conserved in the long terminal repeats (LTRs) of HERV-K-10 and -K-11, and validated the functionality of this site using reporter assays. In summary, we identified RBPs as potential regulators of ERV function and demonstrate a new role for RBM4 in controlling ERV expression.

**Significance Statement:** The expression of endogenous retroviruses (ERVs) appears to have broad impact on human biology. Nevertheless, only a handful of transcriptional regulators of ERV expression are known and to our knowledge no RNA-binding proteins (RBPs) were yet implicated in positive or negative post-transcriptional regulation of ERVs. We implemented a computational pipeline that allowed us to identify RBPs that modulate ERV expression levels. Experimental validation of one of the prime candidates we identified, RBM4, showed that it indeed bound RNAs made from ERVs and negatively regulated the levels of those RNAs. We hereby identify a new layer of ERV regulation by RBPs.

## Introduction

Human endogenous retroviruses (HERVs) are the remnants of several ancient retroviral germline infections that have accumulated over the past 60 million years (1). Today, they account for approximately 8% of our genome (2). Consequently, as a class, ERVs are perpetual members of our virome, the compendium of viruses that exist in or on an organism (3). As life-long residents of all human cells, these sequences of viral origin likely continue to have a considerable impact on our biology, yet they are not fully understood. Many of the integrated viral DNA sequences contain transcription-factor (TF) binding sites and have been exapted as genomic coordinators of a wide range of important processes, including the expression of interferon gamma (IFNγ) responsive transcripts (4). At the transcriptomic level, HERVs seem largely silent in normal tissue, but they are highly active in embryonic stem cells and the earliest stages of development, where they seem indispensable (5–10). HERV expression is also elevated in a wide range of diseases, among others systemic lupus erythematosus (SLE) (11, 12), various cancers (13–15), and multiple sclerosis (16).

Despite their strong association with fundamental biological processes, little is known about if and how these integral parts of our genome and “virome” are regulated to avoid a potential threat to genome and transcriptome integrity (17). Recently, epigenetic silencing by TRIM28 and HDAC1 to suppress ERV expression at the transcriptional level was identified as a crucial mechanism to control HERVs (18–21). Even though to our knowledge, no RNA binding protein (RBP) was identified to interact with HERVs, we hypothesized that HERVs could also be subject to post-transcriptional regulation. The human genome encodes around 1,500 RBPs that bind and regulate localization, stability, adenylation and splicing patterns of coding and non-coding RNAs in a sequence and/or structure dependent manner (22). A recent study showed that many RBPs bind and repress evolutionarily young mobile repetitive elements, including intronic long interspersed nuclear elements (LINEs) to prevent them from interfering with cellular gene expression (23). By analogy, RBPs seem like attractive candidates for regulation of HERV transcripts, but to date, no examples of such interactions have been documented (24).

Until recently, systematic studies of expression patterns of HERVs have been hampered by the lack of adequate bioinformatic tools. In 2015, the Hammell group released TEToolkit (25), which enables family-wise quantification of HERVs and other classes of transposable elements (TEs) from RNA sequencing (RNA-seq) data. In 2018, the Iwasaki and Vincent groups released ERVmap and hervQuant respectively (26, 27), which enable high resolution mapping of sequencing reads to a curated set of approximately 3,200 HERV proviral loci. These pioneering resources enabled us to develop a hybrid computational pipeline to test the hypothesis that RBPs act as regulators of HERV expression. Running our pipeline on two different single-cell and one bulk RNA-seq datasets from the public domain yielded lists of RBPs with correlated or anti-correlated expression to that of HERVs, which represent possible positive regulators or suppressors of HERV function. Among the top-candidates for HERV-repressing RBPs, we identified RNA-Binding Protein Motif 4 (RBM4). Follow-up experiments in a human cell line demonstrated that RBM4 directly bound proviral transcripts of members of the HERV-K and HERV-H families and that loss of RBM4 resulted in an increased abundance of these HERV transcripts, as well as increased expression of its envelope (env) protein. In summary, we provide the first evidence that RBPs can act as post-transcriptional suppressors of ERVs and identify a novel role for the nuclear RBP RBM4 in regulating HERV-K and -H proviruses.

## Results

To our knowledge, no RBP has previously been reported to act directly on HERV transcripts. We hypothesized that they represent prime candidates to regulate HERVs post-transcriptionally. To identify which ones might, we developed a computational pipeline to systematically analyze the HERV transcriptome in RNA-seq data and compare their expression to those of protein coding genes, including a set of 1,542 RBPs (22). First, we quantify expression of all annotated mRNAs and HERVs. Then we use a robustly expressed subset of HERVs and mRNAs to calculate all pairwise correlations and finally aggregate those to find genes that exhibit a correlated or anti-correlated expression pattern to HERVs (Fig. 1A).

**Figure 1.**
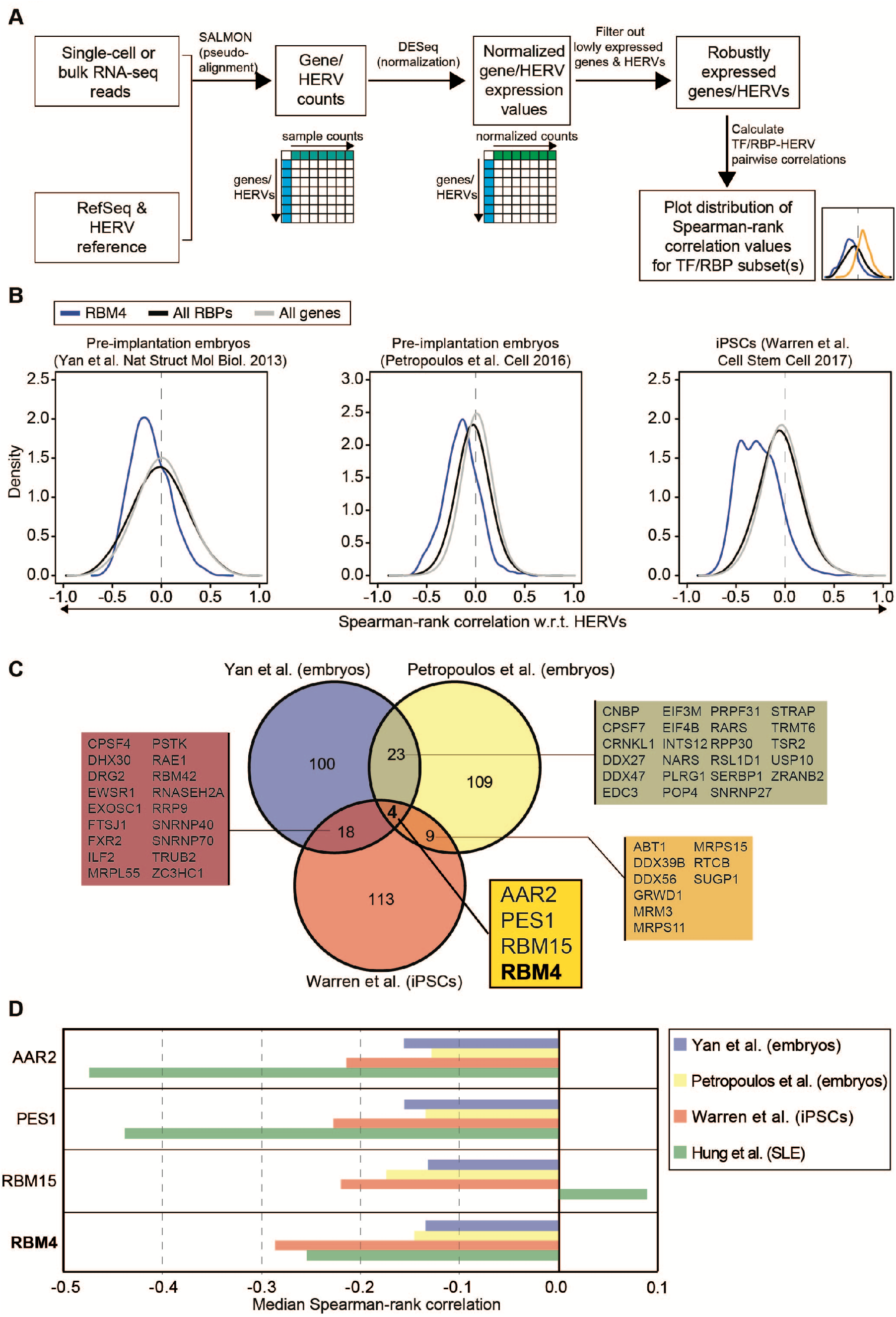
RBM4 expression is inversely correlated with HERVs in multiple transcriptomic datasets from physiologically and pathologically relevant conditions. A) Flow chart of the computational pipeline that was run for each dataset, and used for aligning, quantitating, normalizing and calculating correlations between HERVs versus the expression of mRNAs (including but not limited to those encoding RNA-binding proteins or transcription factors). B) Histograms depict the distribution of Spearman-rank correlation values of RBM4 (blue), all expressed RNA-binding proteins (black) or all expressed genes (gray) versus HERVs in the respective dataset. C) Venn diagram depicts the overlap of the top HERV anti-correlated RBPs (top 10%) that are robustly expressed (> 25th percentile) across the pre-implantation embryos (Yan et al. in blue; Petropoulos et al. in yellow) or iPSC (Warren et al. in red) datasets that were used for panel B. Overlap counts are shown and RBP gene names are listed in the boxes that are matched to the color of the respective overlap region. D) Bar plots depict the median Spearman-rank correlation values for each of the top four RBPs that were consistently anti-correlated across the pre-implantation embryo (Yan *et al*. in blue; Petropoulos *et al*. in yellow) or iPSC datasets (Warren *et al*. in red). The median Spearman-rank correlation values against Hung *et al*.’s SLE dataset are also plotted below the others in green for each of the RBPs.

As proof-of-principle for our pipeline, we sought to rediscover TRIM28 and HDAC1 as suppressors of ERV transcription (18–21), as well as the observed largescale HERV upregulation in SLE patients (26) using an independent dataset of RNA-seqs from peripheral blood mononuclear cells (PBMCs) of 99 SLE patients (28). We found a robust expression of 14,997 RefSeq genes and 1,946 HERVs and the aggregate pairwise correlation of all genes and HERVs indeed showed anti-correlations between HERV expression and epigenetic silencers, including but not limited to TRIM28 and HDAC1 (Fig. S1A, Table S1).

Confident that our pipeline worked as expected, we next focused our attention on finding candidate RBPs that could modulate HERV expression. For our study, we selected three publicly available datasets from independent sources: two single-cell RNA-Seq (scRNA-Seq) datasets of pre-implantation embryos (29, 30) and bulk RNA-Seq of 68 induced pluripotent stem cells (iPSCs) (31). These samples are of particular interest because HERVs are dynamically expressed in pluripotent stem cells and during the earliest stages of development (5–10). Thus, we sought to take advantage of the single-cell variation (n=123) within the former, and the inter-individual variation (n=68) within the latter. Since we were particularly interested in post-transcriptional silencers, we looked for RBPs whose expression anti-correlated with HERVs (Table S2). Four RBPs were consistently found in all datasets among the top 10% anti-correlated RBPs: AAR2, PES1, RBM15, and RBM4 (Fig. 1C,D). Further analysis of the SLE PBMCs also resulted in a negative correlation of RBM4 versus RBPs (Fig. 1D, S1B), although it was not among the top 10% in that dataset (Table S2). Fortuitously, we already had RBM4-deficient human cell lines (Fig. S1C) and some RNA-Seq data from our previous work for immediate analyses (32). We used the popular near haploid cell line called HAP1 (33, 34). This cell line expresses both RBM4 and HERVs, and thus allowed us to dissect the role of this RBP in HERV transcript binding and post-transcriptional silencing.

Analysis of our previous (32) and two new rounds of RNA-Seq data (Table S3) comparing RBM4-deficient cells versus wildtype (WT) controls supported our hypothesis that RBM4 silences HERVs. In the absence of RBM4, a range of HERV transcripts were differentially expressed in the HAP1 model system (Fig. 2A, Table S4). Although, expression of some HERVs were decreased in RBM4-deficient cells, they were either of lower statistical confidence or of much lower baseline abundance. As one example, in the absence of RBM4, reads mapping to ERVmap ID# 4452, a HERV-K-13 (HML4) family member, was dramatically decreased in abundance. (Fig. 2B). However, its baseline expression is around ten-fold lower level than that of the HERV-K (HML2) members (K-10, 6272, K-9, K-21, K-11, 1045, K-7) that were all differentially upregulated (Table S4). Interestingly, there are also HERVs that are expressed but not significantly affected by RBM4 including, but not limited to, K-14 (Fig. 2C). K-14 is interesting for a reason to be discussed later.

**Figure 2.**
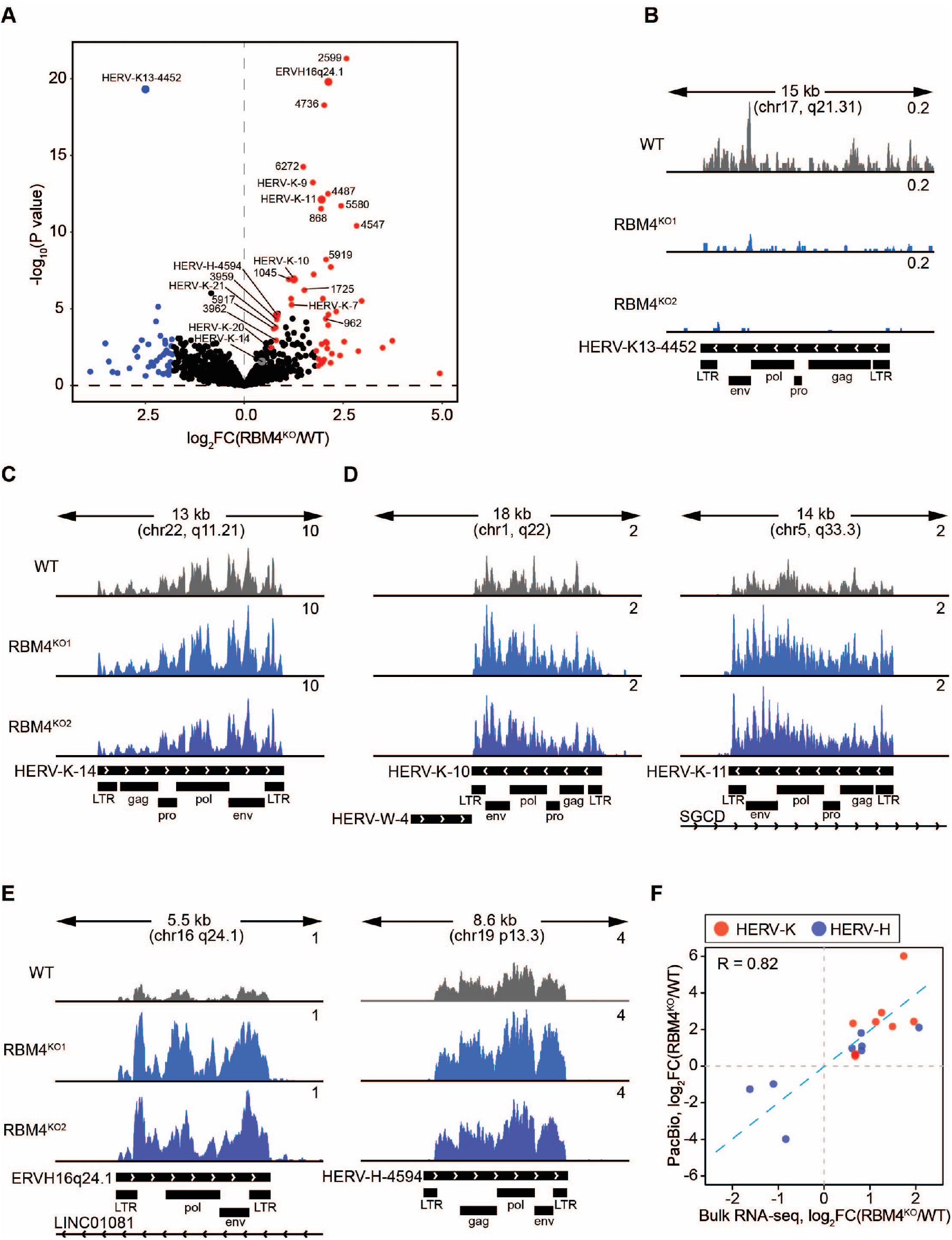
RBM4-deficiency in HAP1 cells leads to increased steady state levels of specific HERV-K and HERV-H transcripts. A) Volcano plot shows differentially expressed HERVs when comparing bulk RNA-seq of RBM4^KO^ HAP1 (clone KO^1^, n=3; clone KO^2^, n=2) versus WT HAP1 (n=3) samples. Highlighted points indicate either HERV transcripts increased (log_2_FC > 2, red) or decreased (log_2_FC < 2, blue) in RBM4^KO^. HERV-H and HERV-K transcripts that are differentially expressed (FDR < 0.05 and baseMean expression > 100) are also colored and labeled with their HERVquant or ERVmap ID. FC = fold change. B) Genome browser shots depict normalized WT or RBM4^KO^ RNA-seq coverage tracks over the decreased HERV-K13 member 4452. The top range of the y-axis (reads per million, RPM) is indicated for each track. The window length and chromosomal location are indicated above the tracks, while the HERV transcript direction (arrow heads) and domains are indicated below the tracks, as well as overlapping parts of annotated RefSeq transcripts. C) Same as B), but for the unchanged HERV-K member, K-14. D) Same as B), but for the increased HERV-K members, K-10 and K-11. E) Same as B), but for the increased HERV-H members, 4417 (aka ERVH16q24.1) and 4594. F) Scatterplot of log_2_FC(RBM4^KO^/WT) values for HERV-Ks (red) and HERV-Hs (blue) that were differentially expressed (FDR < 0.1) in Illumina-based RNA-seq and detected (≥1 aligned read) in PacBio. R = Spearman-rank correlation coefficient.

Overall, more HERV transcripts were elevated in the absence of RBM4 and thus we focused on them. By cross referencing the significantly changed HERVs to the TEToolkit’s (25) repeat-masker annotations, we realized that most HERVs negatively regulated by RBM4 belonged to the HERV-H and HERV-K (HML2) families (35). For instance, at average expression (baseMean value in DESeq2 (36)) >100 and false discovery rate (FDR) <0.05, there are 20 upregulated HERVs in RBM4-deficient samples; 12 of these belong to the HERV-H, and seven belong to the HERV-K (HML2) family. Full analysis results, along with the coordinates and class of the flanking LTRs for all examined HERVs can be found in (Table S4). Illustrative examples of HERV-K (HML2) and HERV-H proviral loci that appeared to be negatively regulated by RBM4 are shown in (Fig. 2D) and (Fig. 2E), respectively. HERV-K (HML2) and HERV-H families represent the evolutionary youngest and transcriptionally most active families of HERVs (37, 38). A major difference between the younger HML2 proviruses and older HML4 proviruses that integrated <10 and >40 M years ago (39), respectively, are their long terminal repeat (LTR) sequences. HML2 proviruses are characterized by LTRs of the LTR5 class, while HML1/4 utilize LTRs of the LTR13 class (35), suggesting that specific sequence elements could mediate the RBM4-effect (see below). Thus, it appears that RBM4 preferentially targets younger mobile elements in analogy to RBPs, such as MATR3 that target younger retrotransposons (23). Of note, changes in HERV expression directly dependent on RBM4 appear to be completely post-transcriptional, since HERVs that were not expressed in WT HAP1 cells were not upregulated in the absence of RBM4. For example, the HERV-W fragment integrated right next to HERV-K-10 is not expressed in neither WT nor RBM4 knock-out (KO) cells (Fig. 2D).

Since short reads from Illumina RNA-seq data are challenging to map to repetitive elements (25), we wanted to confirm the above findings using an orthogonal method, namely, using the long read technology of Pacific Biosciences (PacBio) full-length transcriptome sequencing. This dataset provided around 1.1 (WT) and 1.3 (RBM4 KO) million sequence reads of 2.6 kb average length mapping to the human genome (Table S5). Indeed, the abundances of HERV-K and -H transcripts that could be detected by PacBio correlated well with the quantifications from Illumina RNA-seq data (Fig. 2F).

Next, we wanted to know whether the RBM4-dependent phenotype was specific to HERVs or extended to other types of repetitive elements. We did not observe significant changes in LINE, short interspersed nuclear elements (SINE), or other transposable elements, further suggesting sequence-specificity in RBM4-mediated regulation. HERV-K and -H are clearly the top targets of RBM4-mediated regulation (Fig. 3A, Table S6). To further confirm this finding, we performed reverse transcription and quantitative polymerase chain reactions (RT-qPCR) for three different HERV-K elements: LTR, env, and group antigens (gag). RNA expression of these sequences were consistently elevated in RBM4-deficient cells (Fig. 3B). Finally, these increased RNA levels of HERV-K elements in RBM4 KO cells translated to increased expression of one gene product. Western blotting revealed an increased expression of HERV-K env protein in the absence of RBM4 (Fig. 3C).

**Figure 3.**
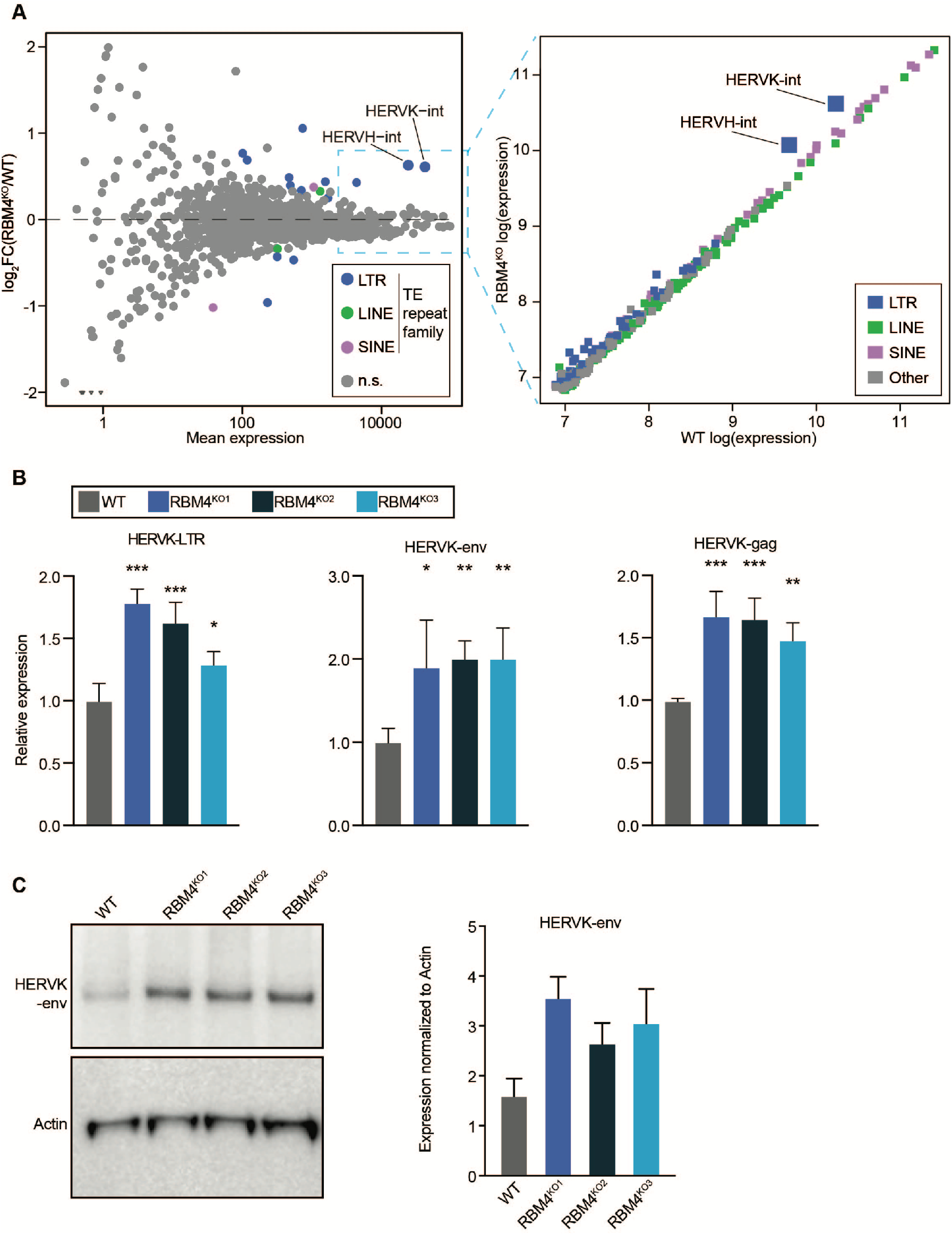
RBM4 deficiency results in increased HERV-K RNA and protein. A) *Left panel*: Mean-difference plot showing differentially expressed TE repeat families. Significantly differentially expressed (FDR < 0.1) families are colored in either blue (LTR family), green (LINE family) or purple (SINE family). Gray (not significant, n.s.) indicates families that are not differentially expressed. The HERV-K and HERV-H families are labeled. *Right panel*: Scatterplot of WT versus RBM4^KO^ log-normalized expression for the 250 most highly expressed TE families. Families are colored by TE family type: LTR (blue), LINE (green), SINE (purple) or other (gray). B) RT-qPCR analysis for HAP1 WT, and three RBM4^KO^ cell lines for the indicated individual HERV-K domain transcripts from independent cultures of HAP1 clones (n=3 for each sample). Values represent expression relative to WT after normalization to the geometric mean of four housekeeping genes, *HPRT1*, *ACTB*, *GAPDH* and *RNA18S* encoding 18S ribosomal RNA (2^−ΔΔCt^). For each transcript, * *P* ≤ 0.05, ** *P* ≤ 0.01 and *** *P* ≤ 0.001 indicate results of *post-hoc* Dunnett’s multiple comparisons test after performing one-way ANOVA on the ΔCt (Ct_housekeeping_ -Ct_target_) values. Error bars are mean ± SD. C) Western Blot analysis and quantitation of HAP1 WT and 3 RBM4^KO^ clones. Images shown are representative of two independent replicates. Blots were probed with anti-HERV-K env antibodies and anti-Actin HRP linked antibodies followed by visualization using chemiluminescence-based imaging. Error bars are mean ± SD for quantification for Western blot analysis.

We hypothesized that RBM4 regulates HERVs by directly binding to their transcripts. To test this possibility, we performed photoactivatable ribonucleoside-enhanced crosslinking and immunoprecipitation (PAR-CLIP) (40). Although, there was a previously published RBM4 PAR-CLIP (41), their deposited data did not contain sequence reads mapping to the human genome (M.H., unpublished) and the authors did not provide us with their raw data upon request. Thus, we performed four independent PAR-CLIP replicates of our own using HAP1 cells stably expressing a FLAG-tagged RBM4 (FLAG-RBM4) transgene under control of a doxycycline-inducible promoter. Following metabolic labeling with 4-thiouridine (4SU) and crosslinking with ultraviolet light (UV) of 312 nm wavelength, we isolated RNA fragments covalently linked to FLAG-RBM4 (Fig. S2A). The RNA recovered from four biological replicates were converted into cDNA libraries and deep sequenced (Table S7). Sequence reads were aligned to the human genome while allowing for multi-mapping in order to account for the repetitive sequence composition of HERVs. The coverage of these aligned reads was evenly distributed across all multi-mapped positions that were of equal mapping quality (see Methods). These distributed reads were then grouped into clusters we consider as probable RBM4 binding sites (Table S7) by PARalyzer (42) to identify those enriched for the diagnostic T-to-C mutations which represent a hallmark of 4SU-mediated crosslinking. On average, we identified 35,933 binding sites, of which ~50% were derived from mRNA intronic regions, consistent with the known nuclear localization of RBM4 protein (Fig. S2B). When we looked at the overlap of annotated targets, we found that 3435 transcripts were reliably bound across all four replicates including RBM4 itself, HERV-K LTR and HERV-H internal (int) sequences (Fig. S2C)

Motif analysis of the clusters using EDlogo (43) and HOMER (44) revealed that RBM4 binding sites were strongly enriched for CGG trimers found in the context of a pyrimidine-rich sequence (Fig. S2D,E). Interestingly, as mentioned above, we identified strong binding of RBM4 to its own mRNA, specifically at exon 3 (Fig. S3A) which suggest the possibility of auto-regulation reminiscent of master regulators such as LIN28B (45–47). Previously, we discovered that RBM4 regulated ULBP1 via alternative splicing (32). Indeed, we found that RBM4 binds around the relevant splice site of ULBP1 (Fig. S3B).

To address our original hypothesis, we mapped the PAR-CLIP reads to a consensus sequence of HERV-H and observed peaks of RBM4-binding across it (Fig. 4A). Then, we mapped the PAR-CLIP reads to a consensus sequence of HERV-K (Fig. 4B) which was feasible since they are over 87% identical in sequence (37). This revealed that RBM4 binding seems to be concentrated at the 5'LTR and 3'LTR of HERV-K while there was less binding to the internal sequences. Since HERV-K-10 and -K-11 transcripts increased in abundance in RBM4 KO cells (Fig. 2D), we inspected the sequences of their LTRs around the peak of RBM4 binding that was most consistent across the four PAR-CLIP replicates (Fig. 4B, lower inset). Perhaps not surprisingly, this region of the LTR is 100% identical between HERV-K-10 and -K-11 since both behave similarly in terms of RBM4-dependency. In contrast, HERV-K-14, which is not regulated by RBM4 (Fig. 2A,C), harbors two natural sequence variants (highlighted as red text in Fig. 4B, lower inset).

**Figure 4.**
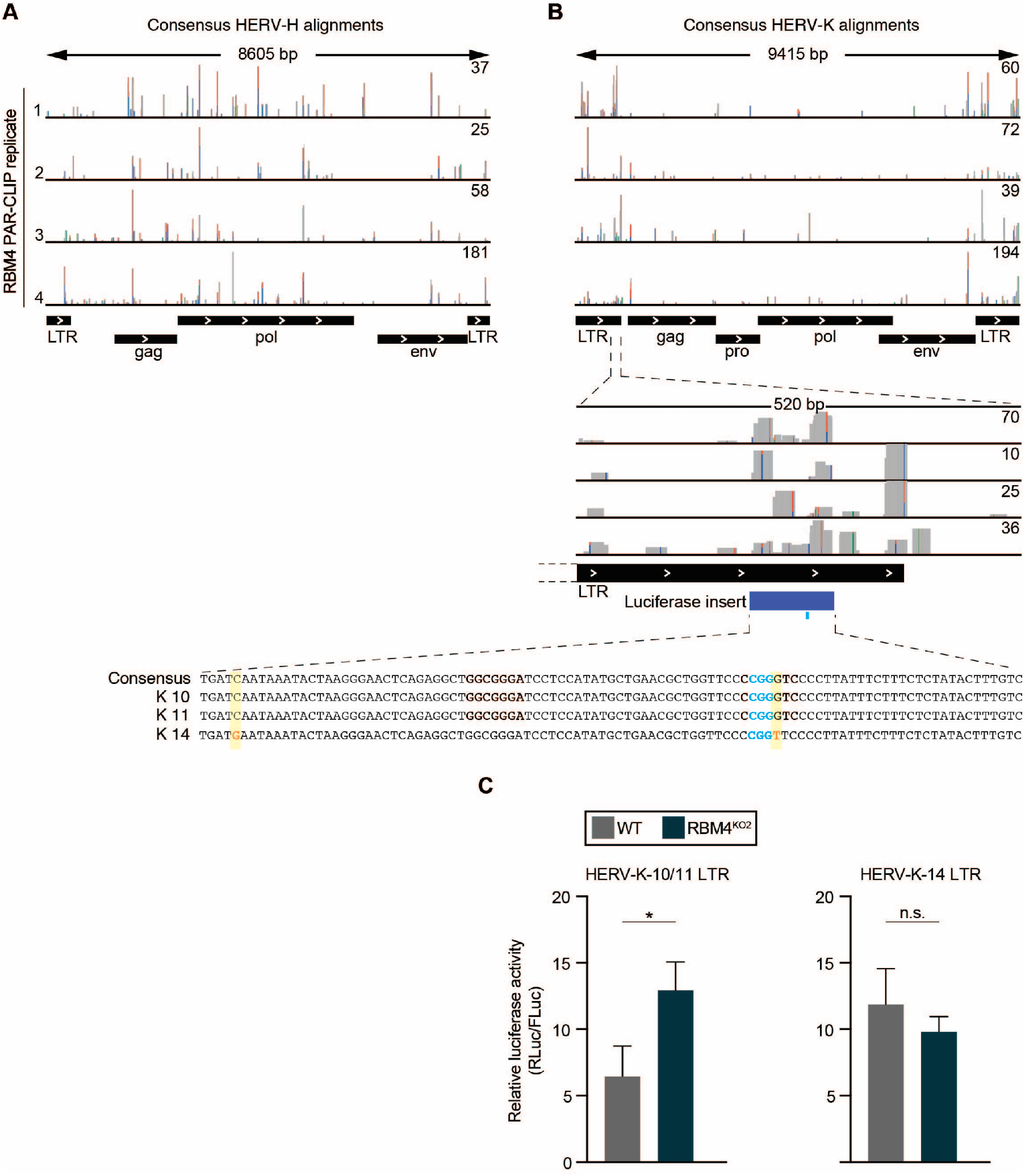
RBM4 binds directly to HERV-K and HERV-H transcripts and regulates their expression. A) Genome browser screenshot shows the coverage of PAR-CLIP reads when aligned to the HERV-H internal consensus sequence from Dfam (DF0000183.4) inside of flanking LTRs (DF0000571.4). The top range of the y-axis (read pileup count) is indicated for each track. Transcript domains are indicated below the tracks when annotations are available. B) *Top panel*: Genome browser screenshot shows as in A) but for the coverage of RBM4 PAR-CLIP reads when aligned to the HERV-K internal (int) consensus sequence from Dfam (DF0000188.4) inside of flanking LTRs (DF0000558.4). *Middle and bottom panels*: Zoomed-in segment of the first LTR shows a validated consensus binding site aligned to the corresponding LTR sequences for HERV-K-10, -K-11 and -K-14. The determinant RBM4 binding motif is shown in blue font, and polymorphisms in HERV-K-14 are highlighted in yellow and red font. C) Ratio of Renilla/Firefly luminescence for HAP1 WT and RBM4^KO2^ cell lines. Cells were transiently nucleofected with reporter constructs containing the HERV-K-10/-11 or -K-14 LTR sequence. * *P* ≤ 0.05 indicate results of two-tailed, independent samples t-test. n.s., not significant. Error bars are mean ± SD.

We decided to take advantage of this experiment of nature and cloned this region into a dual luciferase reporter plasmid. The reporter of HERV-K-10/-11 LTR is highly expressed in the RBM4^KO2^ cell line when compared to WT control (Fig. 4C, left panel). In contrast, the HERV-K-14 LTR reporter is not regulated by RBM4 (Fig. 4C, right panel). Further inspection of these sequences revealed that there are two CGG motifs which could serve as candidate RBM4 binding sites. Since the first motif (on the left) is in a region that is 100% conserved between HERV-K-10, -K-11 and -K-14, we surmised that this motif does not explain the differential regulation observed. The second CGG motif (Fig. 4B, lower inset; highlighted as blue text) is adjacent to a G/T sequence polymorphism suggesting that the differential regulation might map to this single nucleotide difference around where RBM4 binds.

## Discussion

In summary, we mined public transcriptomic data from early life stages and induced pluripotent cells to identify possible RBPs that might regulate HERVs post-transcriptionally. Our *in silico* screen yielded an over-abundance of possible candidates (Table S2) that might act as co-factors or independent factors in a context dependent manner, but only a handful of candidates that consistently passed our filters in all four datasets that we analyzed (Fig. 1C,D). We suspect that this work on RBM4 is only the beginning in recognizing a broader set among the >1,500 human RBPs (22), that act on endogenous and exogenous viruses. After RNA is transcribed from DNA, it can undergo many processing and regulatory steps that control its abundance and/or localization within the cell. This additional layer of regulation at the post-transcriptional level allows cells to fine-tune and/or rapidly control the fate of RNA.

RBM4, the candidate RBP that we followed up on harbors two RNA-recognition motifs (RRM1 and 2) and a CCHC-zinc finger. It was first discovered in the fruitfly (48, 49), but is highly conserved evolutionarily and phylogenetically related to Polypyrimidine Tract Binding Protein 1 (PTBP1) (50). RBM4 is expressed in all tissues and all cell types. In addition to evolutionary conservation, ubiquitous expression, and above mentioned apparent auto-regulation, RBM4’s functional importance is further highlighted by the fact that gnomAD v.2.1 (51) -the largest collection of human genome sequences to date containing data from over 141,000 subjects - does not currently contain a single case of homozygous loss of function of RBM4. RBM4 is commonly described as a splicing factor and a tumor suppressor, yet, there are only a few anecdotal examples of RBM4-mediated, consequential alternative splicing (52–55), and conflicting reports on its association with cancer outcomes (56, 57).

Interestingly, based on analysis of massive public datasets, several bioinformatics and data mining tools predict a role for RBM4 in control of viral infections. For example, Archs4, a website predicting gene functions through massive mining of publicly available RNA-seq data from human and mouse (58), predicts “establishment of integrated proviral latency” to be the top function of RBM4 [https://amp.pharm.mssm.edu/archs4/gene/RBM4]. Similarly, the MetaSignature DataBase (59), which is based on multi-cohort transcriptome analysis of transcriptomic data from over 100 diseases, identifies “Respiratory Syncytial Virus”, “Dengue”, and “Meta-Virus” as top diseases with statistically significant and negative effect sizes with respect to RBM4. Furthermore, The Signaling Pathways Project (https://www.signalingpathways.org/index.jsf) lists the IFNγ receptor family as a potent regulator of RBM4 expression (60). In our own previous work (32), we established that RBM4 is necessary for protein expression of ULBP1, an NKGD2 ligand that acts a signal for recognition of virally infected cells by NK cells. All of these lines of evidence, along with our findings in the present manuscript about modulation of young endogenous retroviruses by RBM4, point to a potential, under-appreciated role for RBM4 as a viral restriction factor. As such, we asked if there is something special about the RBM4 motif that we have identified from PAR-CLIP. Intriguingly, preliminary analysis of viral transcripts from the Virus-Host Database (61) suggests that 6-mers containing RBM4 motif are enriched in viral transcripts compared to human ones (Fig. S4A).

While the relation between RBPs (in particular, RBM4) and exogenous retroviruses still remains to be investigated, we believe that we have provided strong evidence for regulation of young HERVs by RBM4, as a first example of its kind. To our knowledge, this is the first report of PAR-CLIP analysis to demonstrate that HERV transcripts are bound by an RBP. The lack of documented binding of RBPs to HERVs might partly stem from the inherent bioinformatics challenges of dealing with repetitive elements and partly from the limited expression of young HERVs (HERV-K/HML2 and HERV-H) in the commonly used model cell lines like K562 (Fig. S4B,4C). Recent computational tools like TEToolkit (25), ERVmap (26), and HERVQuant (27) have substantially ameliorated the problems on the computational front. As far as the experimental side is concerned, we have now identified HAP1 cells as an ideal cell line and model system for investigating post-transcriptional regulation of HERVs by RBPs.

HAP1 cells were originally obtained in an unsuccessful attempt to induce pluripotency in KBM7 cells (a cell line derived from a male chronic myelogenous leukemia patient) by expression of the “Yamanaka” factors OCT4, SOX2, MYC and KLF4, which resulted in loss of hematopoietic markers (33, 34, 62). The high expression levels of young HERVs in HAP1 cell, which exceeds their expression in KBM7 by orders of magnitude (data not shown) hints at the possibility that they are closer to pluripotent stem cells than commonly assumed. Last but not least, we note that HAP1 cells are often used in genome-wide screens to study infections by exogenous viruses (33, 34, 62). If there is indeed a role for HERVs in cellular defense against such external infections (5), the high level of HERV expression in HAP1 cells might be an unrecognized confounding factor that would merit further consideration.

## Materials and Methods

### Computational Identification of HERV-regulating Candidate RBPs

#### Public datasets

Four relatively large datasets from conditions where HERVs had been previously reported to be highly expressed were chosen. For Hung et al. (28), all SLE samples were processed (n=99). For Yan et al. (29), samples from 4-cell, 8-cell and Morula-stage were considered (n=48). For Petropoulos et al. (30), 81 samples from embryo day 3 were processed, however six samples that had less than one million mapped reads after alignment were subsequently excluded from further analyses, leaving n=75 samples. The single-cell datasets were usable because they were essentially full-length coverage low input RNA-seq as opposed to 3' end profiling (eg. 10X Genomics). For Warren et al. (31) all PBMC-derived iPSC samples were chosen (n=68).

#### Quantification and Correlation Analysis

The sequences for Human RefSeq cDNA and flanking 100 bases were downloaded from ENSEMBL BioMart in December 2018 – ENSEMBL gene 94. (41,599 transcripts, thereof 41,368 coding). Additionally, 3,173 Pre-processed proviral HERV-sequences were obtained from HERVquant (27) (https://unclineberger.org/vincent/resources). These two sets of transcripts were concatenated, and a Salmon master index was created for the combined set of 44,772 transcripts K-mers=31, keeping duplicate sequences. All samples were pseudo-aligned by Salmon (version 0.12.0) (63) and RefSeq and HERV transcripts were quantified using this master index, with following parameters (*--validateMappings --useVBOpt --numBootstraps 25*). The resulting expression matrices (raw read-counts) were imported to R and the counts for different isoforms of the same gene were aggregated to a single, gene-level, expression value. Samples with less than 10^6^ mapped reads were excluded from further analysis (this only removed 6 samples from Petropoulos et al., but did not affect any of the other datasets). For each dataset, the expression levels were normalized by DESeq2 (36), and to ensure robust expression, transcripts below the bottom 25% quartile of baseMean values were filtered out from further analysis. For remaining genes and proviruses, pairwise Spearman’s rank correlation was calculated between each gene and each provirus, and the distribution of these correlations were plotted using density function in R. Importantly, in the case of RBM4, qualitatively similar plots were obtained at much higher filtering thresholds (33%, 50%, 66%, 75%). Overall, these results suggested a robust inverse correlation between RBM4 and HERVs across conditions, sequencing platforms (single end versus paired end), and HERV-expression levels, making it an attractive candidate for follow up.

### Cell Lines

The following were obtained from David Raulet (32): WT, three RBM4-deficient clones (RBM4 KO^1-3^), and RBM4 KO^2^ transduced with FLAG-RBM4 driven by a Dox-inducible promoter. The cell lines were cultured in IMDM medium (Quality Biological) supplemented with 10% (v/v) Tet System Approved fetal calf serum (Takara Bio/previously Clontech), 100 U/mL penicillin, 100 μg/mL streptomycin (Pen/Strep) (Thermo Fisher Scientific), at 37°C in an incubator containing 5% CO_2_.

### Western Analyses

Samples were denatured at 95°C for 10 min, resolved on a 4-20% gradient ExpressPlus™ PAGE gels (GenScript) and transferred to PVDF membranes (Thermo Fisher Scientific) with a Novex X Cell Western Blot Semi-dry system (Thermo Fisher Scientific). Blots were blocked with 5% (w/v) milk in Tris Buffer Saline with 0.1% Tween (TBST buffer) for 1 hour at RT with gentle rocking and probed with primary antibodies at 4°C, overnight with gentle rocking. The blots were washed for 5 min with TBST three times. An HRP-conjugated secondary antibody incubation was followed for 1 hour at room temperature and washed again for 5 min with TBST three times. Blots were developed with SuperSignal West Dura Extended Duration Substrate (Thermo Fisher Scientific) and images were acquired using LAS-4000 (Fuji Film) or Azure c500 Imager (Azure Biosystems). Band densities on blots were quantified with ImageJ (NIH) and background from an adjacent blank area of the blot was subtracted. Signal from the protein band of interest was normalized to β-Actin. Antibodies used were HERV-K env (Austral Biological, HERM-1811-5); RBM4, (Abcam, ab130624); β-actin (Sigma, A3254).

### RNA Isolation

Total RNA was isolated from the HAP1 cells (WT and RBM4 KO) using the RNAzol^®^ RT (Molecular Research Center) per manufacturer’s instruction. RNA was quantified with Qubit^®^ RNA HS Assay Kit according to the manufacturer’s recommendation (Thermo Fisher Scientific). RNA integrity was verified with Agilent RNA 6000 Nano kit on the 2100 Bioanalyzer (Agilent Technologies) or HS RNA Kit (15 nt) on the Fragment Analyzer (Agilent Technologies) according to the manufacturers’ protocol.

### RNA-seq Illumina System

RNA-Seq libraries were prepared from 65 ng of total RNA and 1 μL of 1:5000 (v/v) dilution of ERCC ExFold RNA Spike-In Mix 1 (Thermo Fisher Scientific) with TruSeq^®^ Stranded mRNA Library Prep for NeoPrep™ instrument as per the NeoPrep Library Prep System Guide on the instrument (Illumina).

Libraries were diluted 1:5 (v/v) and 1 μL was used to check the expected size (~300 bp) and the purify was verified with the Agilent High Sensitivity DNA Kit as manufacturer’s recommendation (Agilent Technologies). One μL of the diluted libraries were used to quantify as recommended in the Quant-iT PicoGreen dsDNA Assay Kit (Thermo Fisher Scientific) with standard range from 0 to 14 ng/ well. Libraries were normalized with molecular grade water to 2 nM with the above expected size and quantification.

Pooled libraries of 2 nM were denatured and diluted as per the Standard Normalization Method for the NextSeq® System (Illumina). Final loading concentration of 1.8 pM and 1% PhiX (v/v) was used with the NextSeq^®^ 500/550 High Output Kit v2 (150 cycles) (Illumina). Samples were sequenced paired-end with 75 cycles and single index of 6 cycles.

### RNA-seq Computational Analysis

For the Volcano plot, transcripts and HERVs were quantified using the same pipeline as above, and differential expression of HERVs was determined by DESeq2. Since there is an almost one to one match between hervQuant and ERVmap HERV-collections, the more descriptive ERVmap IDs were used for labeling in the plot.

To determine the familywise changes to transposable elements, the TEToolkit (25) pipeline and recommendations was deployed. Briefly, reads were first mapped to genome (hg38) using STAR (version 2.5.2a) (64), allowing for multi-mapping (parameters *--outFilterMultimapNmax 100 -- winAnchorMultimapNmax 100* as recommended in TEToolkit manual) and then the expression levels of all genes and TEs were quantified by TEToolkit with following parameters: The RPM-normalized bigwig files from the above STAR alignment were used for the screenshots in Fig. 2, corroborating the results obtained by Salmon.

### RNA-seq PacBio System

Total RNA was cleaned via Agencourt RNA Clean XP (Beckman Coulter) beads and resuspended in Buffer EB (Qiagen). Quality and quantity was assessed via BioPhotometer (Eppendorf) and Agilent RNA 6000 Pico kit on the 2100 Bioanalyzer (Agilent Technologies). SMRTbell libraries were generated using Pacific Bioscience’s no-size selection Iso-Seq Template Preparation guide (PN 101-070-200 version 04, November 2017) following the manufacturer’s recommendations. Input amounts for generating cDNA (Takara Bio) was 1000 ng and PCR was optimized at 10 cycles for subsequent large-scale PCR. Fraction 1 and 2 of the cleaned-up products were quantified via Qubit (Thermo Fisher Scientific) fluorometric assay and equal molar quantities were combined resulting in over 1 μg cDNA going forward. Final SMRTbell libraries were quantified using Qubit and qualitative and quantitative analyses were performed using BioAnalyzer DNA 12000 kit (Agilent Technologies), and Fragment Analyzer Large-Fragment kit (Agilent Technologies). Pacific Biosciences stand-alone Excel template was followed for primer and polymerase addition/cleanup and sequence data for each library was generated using MagBead loading on three independent SMRTcells on the Sequel (Pacific Biosciences) at 50, 65, and 75 pM respectively. Movie collection time was set at 10 hours with a pre-extension time of 120 minutes.

### RNA-seq PacBio Computational Analysis

All reads were mapped to hg38 using minimap2 (65) using the following parameters: -*ax splice - uf --secondary=no -C5 Homo_sapiens.GRCh38.dna.primary_assembly.fa*

The resulting bam files were then filtered for mapping quality and unique mapping using samtools view -*F 256 -q 20*, and converted to RPM-normalized bigwig files. Next, region-wise coverage was extracted using derfinder (66) from the bigwig files, and regions overlapping HERV-proviruses were determined using Granges (67).

### DNA oligonucleotides

For sequences see Table S8

### RT-qPCR

First-strand cDNA was synthesized from 2 μg of total RNA using the Maxima Reverse Transcriptase (Thermo Fisher Scientific) with random hexamer primers. Real-time PCR was performed using the Power SYBR green (Thermo Fisher Scientific) 2x master mix or Fast Start Universal SYBR Green (Roche) 2x master mix in a 7900HT fast Real-Time PCR system (Thermo Fisher Scientific) or QuantStudio 7 Flex Real-Time PCR System (Thermo Fisher Scientific) using gene-specific primers for *ACTB*, *HPRT1* and *GAPDH* (32), as well as HERV-K (LTR, gag and env) and *RNA18S* (5) (see Table S8 for sequences). The HERV-K LTR primers are specific for LTR5_Hs (the _Hs denotes specificity for *Homo sapiens*).

### PAR-CLIP

PAR-CLIP was performed as described previously (40, 68). Briefly, for each PAR-CLIP experiment, 500-600 x 10^6^ HAP1 RBM4 KO^2^ cells transduced with FLAG-RBM4 grown in 15 cm plates were used for each biological replicate. 24 hours before UV-crosslinking expression of the FLAG-RBM4 transgene was induced by addition of 200 ng/mL doxycycline (Takara) and 16 hours (or 1 hours) before UV-crosslinking 100 μM 4SU was added. After washing with ice-cold PBS cells were crosslinked (irradiation with 312 nm UV light, 5 min) and scraped off the dishes using a rubber policeman. Cells were lyzed in the equivalent of three cell pellet volumes of lysis buffer (50 mM HEPES, pH7.5, 150 mM KCl, 2 mM EDTA, 0.5 mM DTT, 0.5% (v/v) NP-40, protease inhibitors) and cleared by centrifugation at 13,000g. Cell lysates were treated with 1 U/μL RNase T1 (Thermo Scientific) and then immunoprecipitated using anti-Flag magnetic beads (M2, Sigma Aldrich). The beads were taken up in one original bead suspension volume (10 μL of bead suspension per milliliter of lysate) and incubated with lysate in 15-mL centrifugation tubes on a rotating wheel for 1 h at 4°C. Beads were washed three times in 1 mL of immunoprecipitation wash buffer (50 mM HEPES-KOH at pH 7.5, 300 mM KCl, 0.05% (v/v) NP40, 0.5 mM DTT, complete EDTA-free protease inhibitor cocktail, Roche) and resuspended in 1 vol of immunoprecipitation wash buffer. The RNA residing in the immunoprecipitated was further trimmed with 100 U/μL RNase T1 (Thermo Scientific). The beads were then washed three times in 1 mL of high-salt lysis buffer (50 mM HEPES-KOH at pH 7.5, 500 mM KCl, 0.05% (v/v) NP40, 0.5 mM DTT, complete EDTA-free protease inhibitor cocktail, Roche) and resuspended in one bead volume of dephosphorylation buffer.

3’ ends of the RNA fragments were dephosphorylated using 0.5 U μl^−1^ calf intestinal phosphatase (New England Biolabs) for 10 min at 37°C, shaking. RNA was radioactively 5' end-labeled using 1 U μl^−1^ T4 polynucleotide kinase (Thermo Fisher Scientific) and 0.5 μCi μl^−1 32^P-γ-ATP (PerkinElmer) for 30 min at 37°C. Quantitative 5′end phosphorylation of RNA was accomplished by adding ATP (Thermo Fisher Scientific) to a concentration of 100 μM for 5 min.

The protein–RNA complexes were separated by SDS-PAGE and transferred to nitrocellulose, and RNA–protein complexes were visualized by autoradiography. Radioactive bands corresponding to the FLAG-RBM4 ribonucleoprotein complex were recovered, and protein was removed by digestion with 1.2 mg/mL proteinase K (Roche) in proteinase K buffer (50 mM Tris-HCl at pH 7.5, 75 mM NaCl, 6.25 mM EDTA, 1% SDS) for 2 h at 37°C. Next, the RNA was isolated by acidic phenol/chloroform extraction and ethanol precipitation. The recovered RNA was converted into a cDNA library using a standard small RNA library protocol (69) and sequenced on an Illumina HiSeq2500 or 3000 instrument.

### PAR-CLIP computational analysis

The PAR-CLIP sequencing data were processed by the PARpipe pipeline (https://ohlerlab.mdc-berlin.de/software/PARpipe_119/). Adapters were trimmed using Cutadapt 1.8.3 and reads were aligned to the hg38 (GRCh38.p5) genome using Bowtie v1.1.2 allowing for 2 mismatches, and keeping up to 3600 multi-mapping alignments tied for the best alignment score (parameters: *-v 2 -a -m 3600 --best --strata)*. This was done to account for the highly repetitive nature of HERVs among other transposable elements. The coverage of these reads was evenly divided across all genomic positions which were tied for the best alignment score. The identifiers for the alignments of each read to each distinct genomic position were processed using in-house scripts to be unique and passed into PARalyzer (42) for clustering and annotation.

PARalyzer v1.5 was used to process reads into groups and clusters with the following parameters:

MINIMUM_READ_COUNT_PER_GROUP=5,
MINIMUM_READ_COUNT_PER_CLUSTER=5,
MINIMUM_READ_COUNT_FOR_KDE=3,
MINIMUM_CLUSTER_SIZE=11,
MINIMUM_CONVERSION_LOCATIONS_FOR_CLUSTER=2,
MINIMUM_CONVERSION_COUNT_FOR_CLUSTER=2,
MINIMUM_READ_COUNT_FOR_CLUSTER_INCLUSION=1,
MINIMUM_READ_LENGTH=16,
MAXIMUM_NUMBER_OF_NON_CONVERSION_MISMATCHES=2
CUTOFF=0.3

Reads, groups and clusters were annotated to the features in the Gencode v24 annotation file for reference chromosomes, scaffolds, assembly patches and alternative loci, with introns given higher precedence in the annotation rank as compared to coding regions. This resulted in 25,371 clusters for combined +Dox, and 885 clusters for combined −Dox control samples.

### PAR-CLIP Motif Enrichment Analysis

Clusters from +Dox samples overlapping clusters from −Dox samples were excluded from further analysis using bedtools. The resulting, filtered cluster BED file was used as input to HOMER (44) with *-rna* and *-keepFiles* flags. Sequence logos were re-plotted with positional information score (bits) using HOMER’s positional weight matrix (.motif) files as input to the R package, ggseqlogo (https://github.com/omarwagih/ggseqlogo).

An intermediate product of the above HOMER analysis was the extended sequences centered around each cluster. The set of non-redundant extended sequences were used as input to Logolas (43), using the frequencies of nucleotides in human genome (A=0.29,C=0.21,G=0.21,T=0.29) as background.

### Consensus HERV-K and HERV-H Reference Generation and Analysis

The custom consensus genome for HERV-K and -H was generated by directly taking the sequences from the publicly available transposable element database, Dfam 3.1 (70) (https://dfam.org/home). The internal HERV (gag, pro, pol, env and spacers) sequences were inserted in between their respective canonical long terminal repeat (LTR) sequences to form the consensus sequences. Respectively, these are:

HERV-K internal: https://dfam.org/family/DF0000188/summary
HERV-K LTR: https://dfam.org/family/DF0000558/summary

HERV-H internal: https://dfam.org/family/DF0000183/summary
HERV-H LTR: https://dfam.org/family/DF0000571/summary

The reference GTF annotation files were generated using the reported internal HERV domain intervals from the above links. The RBM4 PAR-CLIP libraries described above were then re-run through PARpipe/PARalyzer using this custom HERV-K and HERV-H reference genome and annotation, using default parameters (without any multi-mapping modifications described in the above PAR-CLIP analysis section). The output sequence alignment map (SAM) files were then expanded, filtered for forward (matching) strand, sorted, indexed and then loaded into IGV (Integrative Genomics Viewer) (71) along with the custom reference genome and annotation for visualization.

### Plasmids

To construct luciferase reporter plasmids, we cloned synthetic fragments of HERV DNA (sequences in Table S8) using gBlocks^®^ (Integrated DNA Technologies) into psiCHECK-2 (Promega). The fragments were inserted downstream of the Renilla luciferase gene using XhoI and NotI restriction sites and DNA ligation. Plasmid inserts were verified by sequencing (Macrogen).

### Luciferase Reporter Assay

5×10^5^ HAP1 cells were transiently transfected with 300 ng of plasmid using the SE cell line 4D-Nucleofector kit S (Lonza), on a 4D nucleofection unit (Lonza) using protocol DZ-113. Cells were plated out evenly in three separate wells on a 12-well plate, cultured for 24 hours, and luciferase reporter activity was assessed using Dual Luciferase Reporter Assay System (Promega) according to the manufacturer’s protocol. Luminescence measurements were made in a FLUOstar Omega instrument (BMG LABTECH).

## Acknowledgments

We thank David Raulet (UC Berkeley) for providing RBM4 KO cells; Jingwen Gu and Yunhua Zhu (Bioinformatics and Computational Biosciences Branch, NIAID) for assistance with statistical analyses of RT-qPCR; Gustavo Gutierrez-Cruz and Stefania Dell'Orso (NIAMS) for sequencing PAR-CLIP libraries; NIAID Office of Cyber Infrastructure and Computational Biology for high-performance computing; Wesley Tung and Helen Su (NIAID) for luminometer; members of Muljo group, Astrid Haase (National Institute of Diabetes and Digestive and Kidney Diseases), Chrysi Kanellopoulou (Incyte Research Institute) and Ron Germain (NIAID) for comments and suggestions. This research was supported by the NIH Intramural Research Program of NIAID and NIAMS.

## Supplementary Figures and Legends

**Figure S1.**
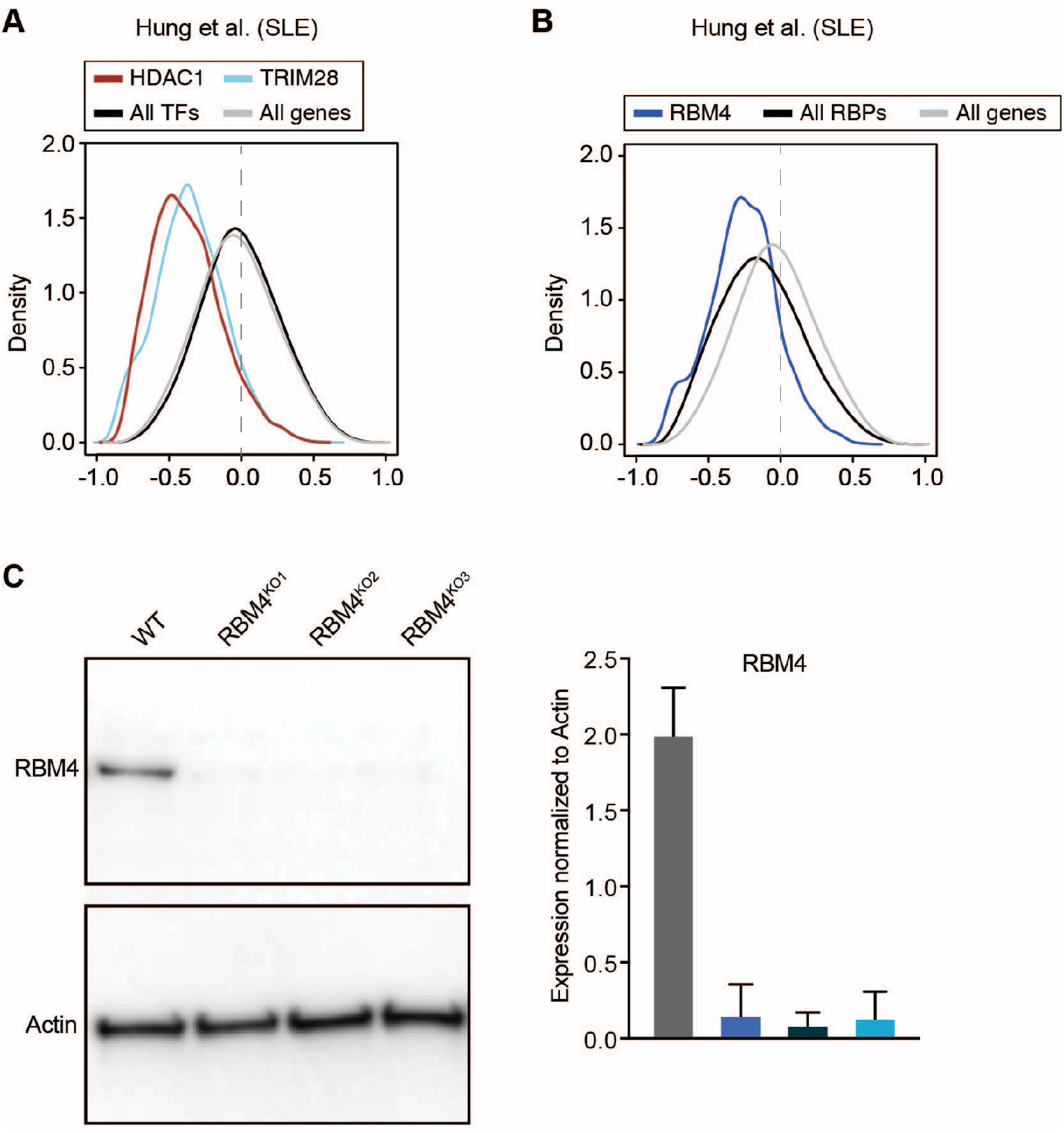
Results from *in silico* screen and Western blot validation of RBM4^KO^ clones. A) Histogram depicts the distribution of Spearman-rank correlation values of the transcriptional regulators, HDAC1 (brown), TRIM28 (light blue), all expressed transcription factors (TFs, black), or all expressed genes (gray) versus HERVs in the indicated dataset. B) Histogram depicts the distribution of Spearman-rank correlation values of RBM4 (blue), all expressed RNA-binding proteins (black), or all expressed genes (gray) versus HERVs in the indicated dataset. C) Western blot analysis and quantitation of HAP1 WT and three independent RBM4^KO^ clones. Images shown are representative of two independent replicates. Blots were probed with anti-RBM4 antibodies and anti-Actin HRP linked antibodies followed by visualization using chemiluminescence-based imaging. Error bars are mean ± SD for quantification for Western blot analysis.

**Figure S2.**
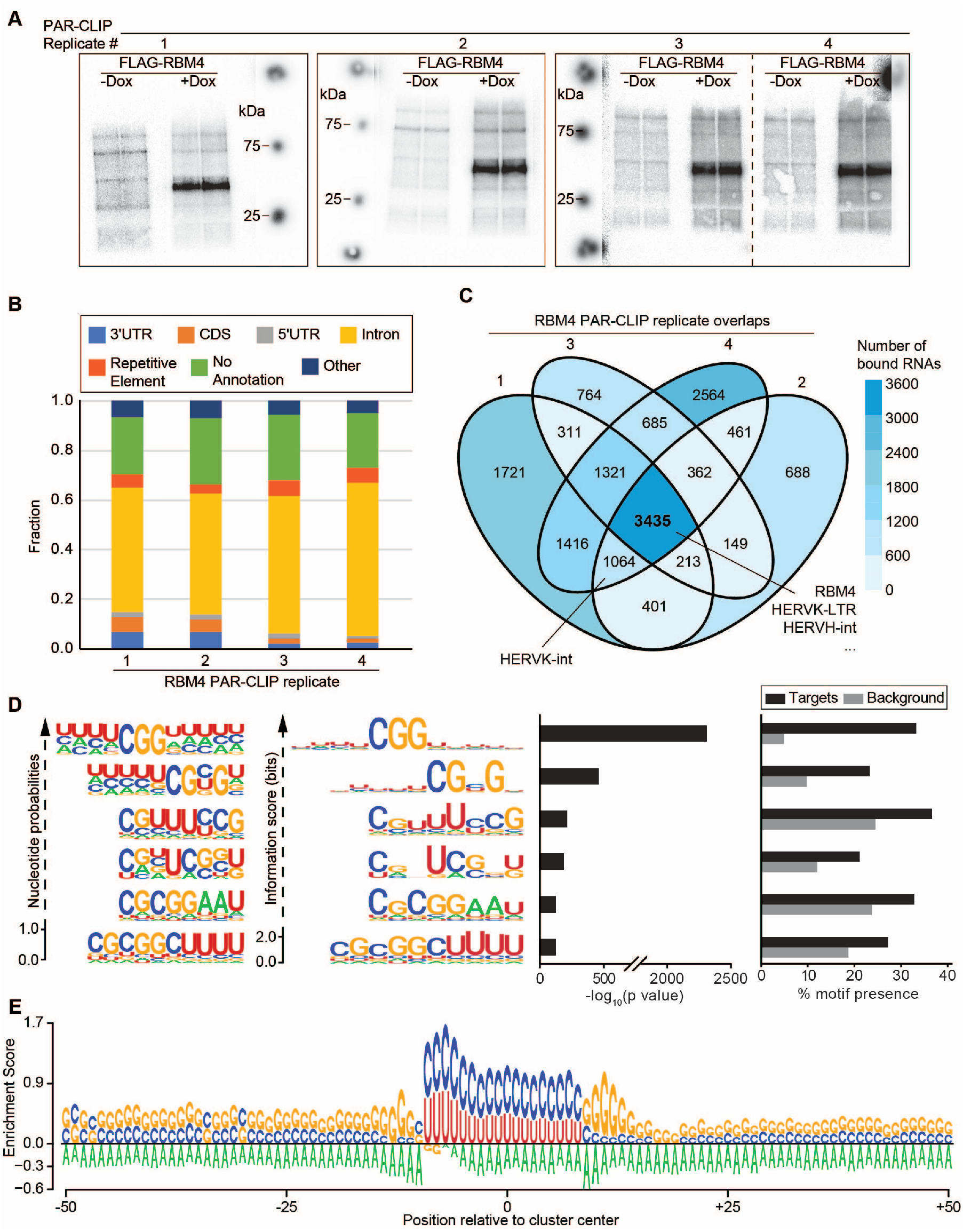
RBM4 PAR-CLIP and motif analyses. A) Autoradiographs of SDS-PAGE fractionating UV-crosslinked, radiolabeled and immunoprecipitated FLAG-RBM4 from cells treated with or without doxycycline (Dox). B) Stacked barplots show the relative fractions of transcriptomic features assigned to PARalyzer-called RBM4 PAR-CLIP clusters for each replicate. C) Venn diagram depicts overlap counts of bound RNA targets across the four RBM4 PAR-CLIP replicates. Overlap sections are colored in light blue to dark blue, with darker blue sections denoting higher overlap counts as indicated in the legend. RBM4, HERV-K-LTR and HERV-H-int are confidently bound in all four replicates, while HERV-K-int is shown to be bound in three out of four. D) HOMER sequence logos and results shown for the top six most significantly enriched RBM4 motifs of nucleotide length 8, 10 or 12. RNA sequence logos in the left panels are plotted with either the positional nucleotide probabilities or information content in bits. Barplots in the right panel show HOMER enrichment statistics by either log-transformed *P* value or relative % presence in target versus background. E) EDLogo plot describing positional nucleotide enrichments in a window ± 50 bp around RBM4 clusters.

**Figure S3.**
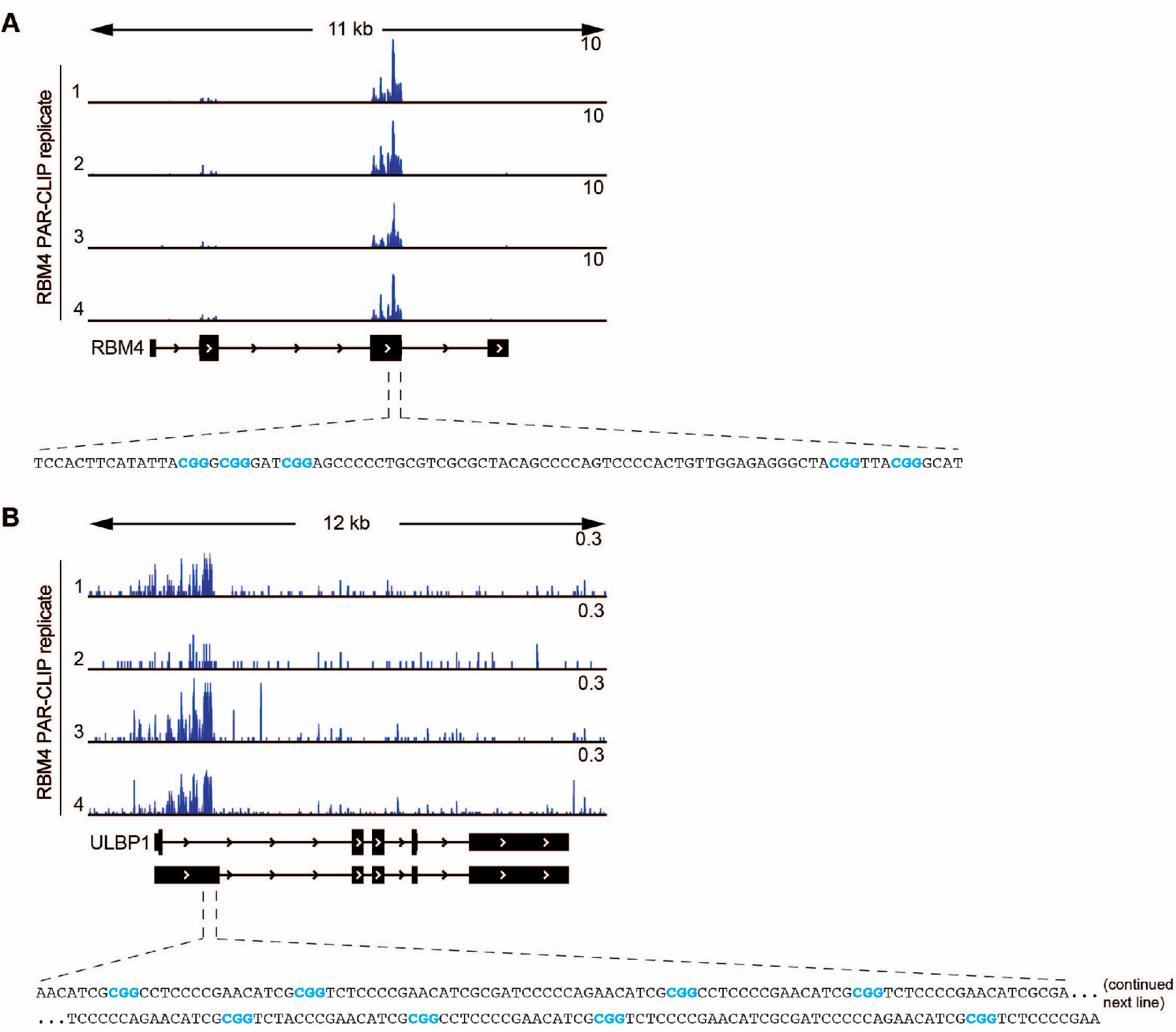
RBM4 binds its own RNA as well as the previously reported ULBP1 alternative splice site. A) Genome browser screenshot depicts normalized RBM4 PAR-CLIP read coverage aligned to the *RBM4* transcript. The top range of the y-axis (reads per million, RPM) is indicated for each track. *Bottom panel*: Zoomed-in segment of the third exon shows the nucleotide sequence of the bound site with instances of the trinucleotide motif, CGG, highlighted in blue font. B) As in A) but for *ULBP1* transcript. *Bottom panel*: Zoomed-in segment near the alternative 5′ splice site shows the nucleotide sequence of the bound site with instances of the trinucleotide motif, CGG, highlighted in blue font.

**Figure S4.**
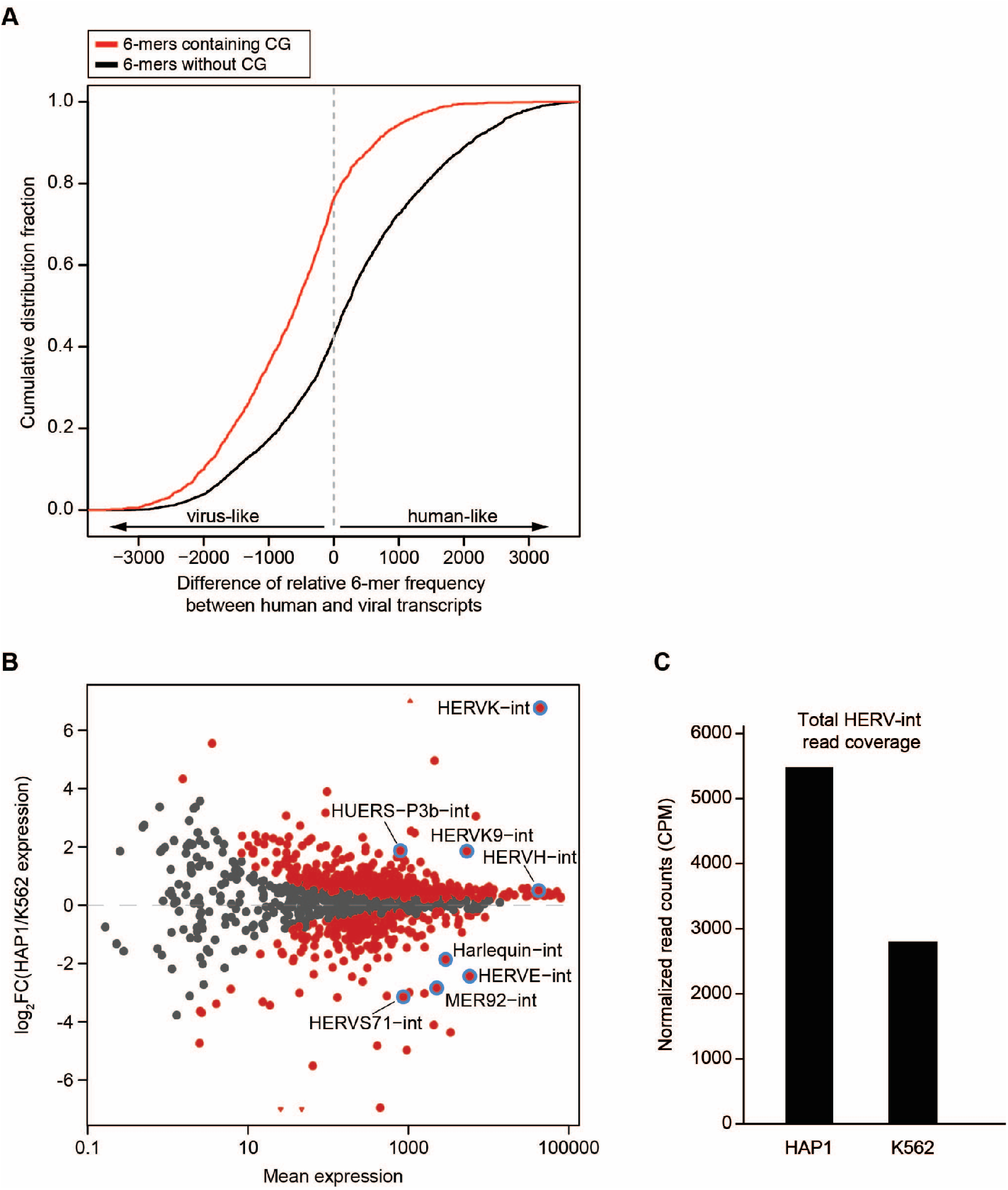
Enrichment of CG-containing hexamers in viral transcriptomes, and comparison of HAP1 versus K562 ERVome. A) The frequency of each 6-mer in all human RefSeq transcripts was determined and these 6-mers were ranked by their frequency, giving us a value *rh* for each 6-mer. We repeated this for all known viral transcripts (from Virus Host Database), determining a value *rv* for each 6-mer. The plot shows the cumulative distribution of the difference for the relative ranks (*rh*-*rv*) of CG-containing 6-mers (red) and of 6-mers lacking CG (black). 6-mers that contain CG (e.g. those compatible with our RBM4 motif) tend to be more frequent in viral transcripts than in human transcripts. B) Mean-difference plot showing TE repeat families differentially expressed (colored in red) between HAP1 and K562 cells. TE repeat families of interest are labeled and circled in blue. C) Plotted are averages of normalized read counts (CPM) for HERV internal sequences as quantified with TEToolkit when re-analyzing Illumina RNA-seq data from WT HAP1 (n=3) or K562 (n=2 from SRA study ERX3372594).

**Table S1.** TF vs HERV Spearman-rank correlation table (see Excel file)

**Table S2.** RBP vs HERV Spearman-rank correlation table (see Excel file)

**Table S3.** Summary statistics for bulk RNA-seq of WT, RBM4^KO1^, and RBM4^KO2^ HAP 1cell lines

| Sample | Total reads | Uniquely mapped reads | Uniquely mapped reads % | Multi-mapped reads | Multi-mapped reads % | Unmapped too short % |
| --- | --- | --- | --- | --- | --- | --- |
| WT 1 | 55805018 | 48541949 | 86.98% | 2391167 | 4.28% | 8.40% |
| WT 2 | 46782145 | 40760526 | 87.13% | 1978451 | 4.23% | 8.36% |
| WT 3 | 54015021 | 46839753 | 86.72% | 2240955 | 4.15% | 8.69% |
| RBM4<sup>KO1</sup> 1 | 43002833 | 36561859 | 85.02% | 1837600 | 4.27% | 10.07% |
| RBM4<sup>KO1</sup> 2 | 69864769 | 61829553 | 88.50% | 2911418 | 4.17% | 6.72% |
| RBM4<sup>KO1</sup> 3 | 51762791 | 45767497 | 88.42% | 2263177 | 4.37% | 6.70% |
| RBM4<sup>KO2</sup> 1 | 52780057 | 45603468 | 83.76% | 2272346 | 4.31% | 8.93% |
| RBM4<sup>KO2</sup> 2 | 60445026 | 53460749 | 88.45% | 2617640 | 4.33% | 6.61% |

**Table S4.** HERVs differentially expressed between HAP1 WT vs RBM4 KO (see Excel file)

**Table S5.** Summary statistics for PacBio long-read RNA-seq of WT and RBM4^KO2^

| Sample | Total reads | Full length nonchimeric reads | Mean Read length (bp) |  | Number of Mapped reads with Flag 256, q 20 |  |  |
| --- | --- | --- | --- | --- | --- | --- | --- |
|  |  |  | All reads | Full length nonchimeric | Total | Full length nonchimeric | Full length nonchimeric + high quality |
| WT | 1314947 | 618045 | 2658 | 3131 | 1144156 | 136087 | 43175 |
| RBM4<sup>KO2</sup> | 1601442 | 657979 | 2634 | 3214 | 1321605 | 126275 | 41558 |

**Table S6.** Differentially expressed HERV families based on TEToolkit (see Excel file)

**Table S7.** Summary results for FLAG-RBM4 PAR-CLIPs in HAP1 cells (see Excel file)

Sheet 1: Summary statistics.

Sheets 2-6: The chromosome, coordinates, IDs, annotated gene, nucleotide sequence, read counts and T2C fraction of binding sites (clusters) obtained from the RBM4 PAR-CLIP, as called by PARalyzer when accounting for read multi-mapping (see Methods). Data for each replicate and the pooled data from all replicates can be found in separate sheets.

**Table S8.**
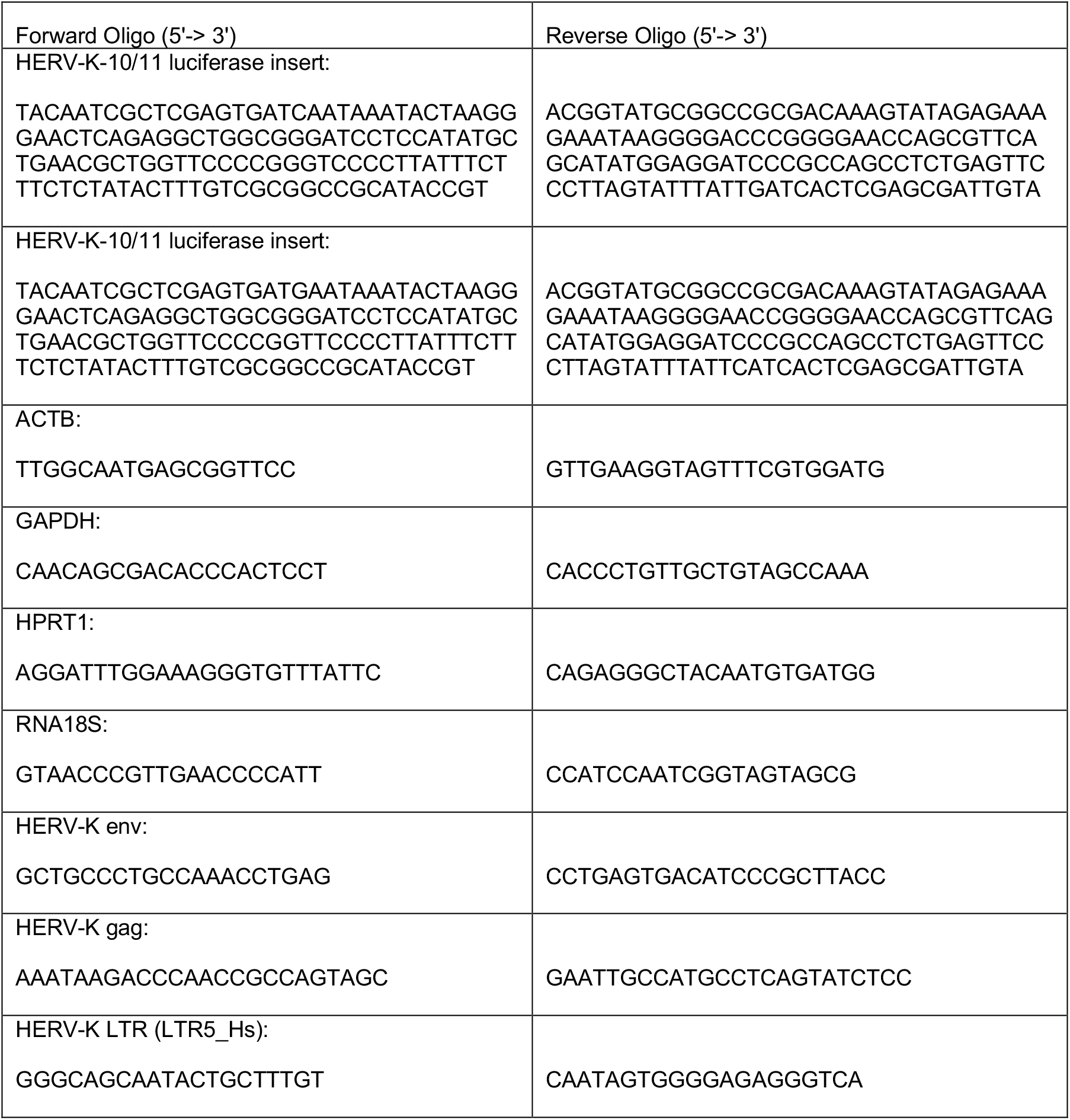
Oligonucleotides used for the luciferase reporter and RT-qPCR experiments.

## References

1. C. Feschotte, C. Gilbert, Endogenous viruses: insights into viral evolution and impact on host biology. Nat Rev Genet 13, 283–296 (2012).

2. E. S. Lander et al., Initial sequencing and analysis of the human genome. Nature 409, 860–921 (2001).

3. H. W. Virgin, E. J. Wherry, R. Ahmed, Redefining chronic viral infection. Cell 138, 30–50 (2009).

4. E. B. Chuong, N. C. Elde, C. Feschotte, Regulatory evolution of innate immunity through co-option of endogenous retroviruses. Science 351, 1083–1087 (2016).

5. E. J. Grow et al., Intrinsic retroviral reactivation in human preimplantation embryos and pluripotent cells. Nature 522, 221–225 (2015).

6. F. A. Santoni, J. Guerra, J. Luban, HERV-H RNA is abundant in human embryonic stem cells and a precise marker for pluripotency. Retrovirology 9, 111 (2012).

7. L. Robbez-Masson, H. M. Rowe, Retrotransposons shape species-specific embryonic stem cell gene expression. Retrovirology 12, 45 (2015).

8. X. Lu et al., The retrovirus HERVH is a long noncoding RNA required for human embryonic stem cell identity. Nat Struct Mol Biol 21, 423–425 (2014).

9. T. W. Theunissen et al., Molecular Criteria for Defining the Naive Human Pluripotent State. Cell Stem Cell 19, 502–515 (2016).

10. J. Goke et al., Dynamic transcription of distinct classes of endogenous retroviral elements marks specific populations of early human embryonic cells. Cell Stem Cell 16, 135–141 (2015).

11. R. S. Treger et al., The Lupus Susceptibility Locus Sgp3 Encodes the Suppressor of Endogenous Retrovirus Expression SNERV. Immunity 50, 334–347 e339 (2019).

12. A. Perl, D. Fernandez, T. Telarico, P. E. Phillips, Endogenous retroviral pathogenesis in lupus. Curr Opin Rheumatol 22, 483–492 (2010).

13. K. Buscher et al., Expression of human endogenous retrovirus K in melanomas and melanoma cell lines. Cancer Res 65, 4172–4180 (2005).

14. F. Wang-Johanning et al., Expression of human endogenous retrovirus k envelope transcripts in human breast cancer. Clin Cancer Res 7, 1553–1560 (2001).

15. D. Diaz-Carballo et al., Cytotoxic stress induces transfer of mitochondria-associated human endogenous retroviral RNA and proteins between cancer cells. Oncotarget 8, 95945–95964 (2017).

16. K. Schmitt et al., Comprehensive analysis of human endogenous retrovirus group HERV-W locus transcription in multiple sclerosis brain lesions by high-throughput amplicon sequencing. J Virol 87, 13837–13852 (2013).

17. R. L. Cosby, N. C. Chang, C. Feschotte, Host-transposon interactions: conflict, cooperation, and cooption. Genes Dev 33, 1098–1116 (2019).

18. T. H. Kao et al., Ectopic DNMT3L triggers assembly of a repressive complex for retroviral silencing in somatic cells. J Virol 88, 10680–10695 (2014).

19. G. Wolf, D. Greenberg, T. S. Macfarlan, Spotting the enemy within: Targeted silencing of foreign DNA in mammalian genomes by the Kruppel-associated box zinc finger protein family. Mob DNA 6, 17 (2015).

20. S. Schlesinger, S. P. Goff, Retroviral transcriptional regulation and embryonic stem cells: war and peace. Mol Cell Biol 35, 770–777 (2015).

21. M. Tokuyama et al., Antibody against envelope protein from human endogenous retrovirus activates neutrophils in systemic lupus erythematosus. bioRxiv 10.1101/776468, 776468 (2019).

22. S. Gerstberger, M. Hafner, T. Tuschl, A census of human RNA-binding proteins. Nat Rev Genet 15, 829–845 (2014).

23. J. Attig et al., Heteromeric RNP Assembly at LINEs Controls Lineage-Specific RNA Processing. Cell 174, 1067–1081.e1017 (2018).

24. J. Attig, J. Ule, Genomic Accumulation of Retrotransposons Was Facilitated by Repressive RNA-Binding Proteins: A Hypothesis. Bioessays 41, e1800132 (2019).

25. Y. Jin, O. H. Tam, E. Paniagua, M. Hammell, TEtranscripts: a package for including transposable elements in differential expression analysis of RNA-seq datasets. Bioinformatics 31, 3593–3599 (2015).

26. M. Tokuyama et al., ERVmap analysis reveals genome-wide transcription of human endogenous retroviruses. Proceedings of the National Academy of Sciences 115, 201814589 (2018).

27. C. C. Smith et al., Endogenous retroviral signatures predict immunotherapy response in clear cell renal cell carcinoma. J Clin Invest 128, 4804–4820 (2018).

28. T. Hung et al., The Ro60 autoantigen binds endogenous retroelements and regulates inflammatory gene expression. Science 350, 455–459 (2015).

29. L. Yan et al., Single-cell RNA-Seq profiling of human preimplantation embryos and embryonic stem cells. Nature structural & molecular biology 20, 1131–1139 (2013).

30. S. Petropoulos et al., Single-Cell RNA-Seq Reveals Lineage and X Chromosome Dynamics in Human Preimplantation Embryos. Cell 165, 1012–1026 (2016).

31. C. R. Warren et al., Induced Pluripotent Stem Cell Differentiation Enables Functional Validation of GWAS Variants in Metabolic Disease. Cell Stem Cell 20, 547–557.e547 (2017).

32. B. G. Gowen et al., A forward genetic screen reveals novel independent regulators of ULBP1, an activating ligand for natural killer cells. eLife 4, 1876 (2015).

33. J. E. Carette et al., Haploid genetic screens in human cells identify host factors used by pathogens. Science 326, 1231–1235 (2009).

34. J. E. Carette et al., Global gene disruption in human cells to assign genes to phenotypes by deep sequencing. Nat Biotechnol 29, 542–546 (2011).

35. L. Vargiu et al., Classification and characterization of human endogenous retroviruses; mosaic forms are common. Retrovirology 13, 7 (2016).

36. M. I. Love, W. Huber, S. Anders, Moderated estimation of fold change and dispersion for RNA-seq data with DESeq2. Genome Biol 15, 550 (2014).

37. R. P. Subramanian, J. H. Wildschutte, C. Russo, J. M. Coffin, Identification, characterization, and comparative genomic distribution of the HERV-K (HML-2) group of human endogenous retroviruses. Retrovirology 8, 90 (2011).

38. P. Gemmell, J. Hein, A. Katzourakis, The Exaptation of HERV-H: Evolutionary Analyses Reveal the Genomic Features of Highly Transcribed Elements. Front Immunol 10, 1339 (2019).

39. K. Hanke, O. Hohn, N. Bannert, HERV-K(HML-2), a seemingly silent subtenant - but still waters run deep. APMIS 124, 67–87 (2016).

40. M. Hafner et al., Transcriptome-wide identification of RNA-binding protein and microRNA target sites by PAR-CLIP. Cell 141, 129–141 (2010).

41. J. Uniacke et al., An oxygen-regulated switch in the protein synthesis machinery. Nature 486, 126–129 (2012).

42. D. L. Corcoran et al., PARalyzer: definition of RNA binding sites from PAR-CLIP short-read sequence data. Genome Biol 12, R79 (2011).

43. K. K. Dey, D. Xie, M. Stephens, A new sequence logo plot to highlight enrichment and depletion. BMC Bioinformatics 19, 473 (2018).

44. S. Heinz et al., Simple combinations of lineage-determining transcription factors prime cis-regulatory elements required for macrophage and B cell identities. Mol Cell 38, 576–589 (2010).

45. M. L. Wilbert, G. W. Yeo, Genome-wide approaches in the study of microRNA biology. Wiley Interdiscip Rev Syst Biol Med 3, 491–512 (2011).

46. M. Hafner et al., Identification of mRNAs bound and regulated by human LIN28 proteins and molecular requirements for RNA recognition. RNA 19, 613–626 (2013).

47. S. Wang et al., Enhancement of LIN28B-induced hematopoietic reprogramming by IGF2BP3. Genes Dev 33, 1048–1068 (2019).

48. F. R. Jackson, S. Banfi, A. Guffanti, E. Rossi, A novel zinc finger-containing RNA-binding protein conserved from fruitflies to humans. Genomics 41, 444–452 (1997).

49. M. A. Markus, B. J. Morris, Lark is the splicing factor RBM4 and exhibits unique subnuclear localization properties. DNA Cell Biol 25, 457–464 (2006).

50. Y. S. Brooks et al., Functional pre-mRNA trans-splicing of coactivator CoAA and corepressor RBM4 during stem/progenitor cell differentiation. J Biol Chem 284, 18033–18046 (2009).

51. K. J. Karczewski et al., Variation across 141,456 human exomes and genomes reveals the spectrum of loss-of-function intolerance across human protein-coding genes. bioRxiv 10.1101/531210, 531210 (2019).

52. D. D, K. Y. Hung, W. Y. Tarn, RBM4 Modulates Radial Migration via Alternative Splicing of Dab1 during Cortex Development. Mol Cell Biol 38 (2018).

53. J. C. Lin, W. Y. Tarn, Exon selection in alpha-tropomyosin mRNA is regulated by the antagonistic action of RBM4 and PTB. Mol Cell Biol 25, 10111–10121 (2005).

54. J. C. Lin, W. Y. Tarn, RBM4 down-regulates PTB and antagonizes its activity in muscle cell-specific alternative splicing. J Cell Biol 193, 509–520 (2011).

55. C. H. Su, K. Y. Hung, S. C. Hung, W. Y. Tarn, RBM4 Regulates Neuronal Differentiation of Mesenchymal Stem Cells by Modulating Alternative Splicing of Pyruvate Kinase M. Mol Cell Biol 37 (2017).

56. Y. Wang et al., The splicing factor RBM4 controls apoptosis, proliferation, and migration to suppress tumor progression. Cancer Cell 26, 374–389 (2014).

57. M. A. Markus, Y. H. Yang, B. J. Morris, Transcriptome-wide targets of alternative splicing by RBM4 and possible role in cancer. Genomics 107, 138–144 (2016).

58. A. Lachmann et al., Massive mining of publicly available RNA-seq data from human and mouse. Nat Commun 9, 1366 (2018).

59. W. A. Haynes et al., Integrated molecular, clinical, and ontological analysis identifies overlooked disease relationships. bioRxiv 10.1101/214833, 214833 (2018).

60. L. B. Becnel et al., Discovering relationships between nuclear receptor signaling pathways, genes, and tissues in Transcriptomine. Sci Signal 10 (2017).

61. T. Mihara et al., Linking Virus Genomes with Host Taxonomy. Viruses 8, 66 (2016).

62. J. E. Carette et al., Ebola virus entry requires the cholesterol transporter Niemann-Pick C1. Nature 477, 340–343 (2011).

63. R. Patro, G. Duggal, M. I. Love, R. A. Irizarry, C. Kingsford, Salmon provides fast and bias-aware quantification of transcript expression. Nat Methods 14, 417–419 (2017).

64. A. Dobin et al., STAR: ultrafast universal RNA-seq aligner. Bioinformatics 29, 15–21 (2013).

65. H. Li, Minimap2: pairwise alignment for nucleotide sequences. Bioinformatics 34, 3094–3100 (2018).

66. L. Collado-Torres et al., Flexible expressed region analysis for RNA-seq with derfinder. Nucleic Acids Res 45, e9 (2017).

67. M. Lawrence et al., Software for computing and annotating genomic ranges. PLoS Comput Biol 9, e1003118 (2013).

68. D. Benhalevy, H. L. McFarland, A. A. Sarshad, M. Hafner, PAR-CLIP and streamlined small RNA cDNA library preparation protocol for the identification of RNA binding protein target sites. Methods 118-119, 41–49 (2017).

69. M. Hafner et al., Identification of microRNAs and other small regulatory RNAs using cDNA library sequencing. Methods 44, 3–12 (2008).

70. R. Hubley et al., The Dfam database of repetitive DNA families. Nucleic Acids Res 44, D81–89 (2016).

71. J. T. Robinson et al., Integrative genomics viewer. Nat Biotechnol 29, 24–26 (2011).

